# High-gamma activity is coupled to low-gamma oscillations in precentral cortices and modulates with movement and speech

**DOI:** 10.1101/2023.02.13.528325

**Authors:** Jeffrey Z. Nie, Robert D. Flint, Prashanth Prakash, Jason K. Hsieh, Emily M. Mugler, Matthew C. Tate, Joshua M. Rosenow, Marc W. Slutzky

**Affiliations:** Southern Illinois University School of Medicine, Springfield, IL 62794, USA; Department of Neurology, Northwestern University, Chicago IL 60611, USA; Department of Neurosurgery, Neurological Institute, Cleveland Clinic Foundation, Cleveland, Ohio, USA; Department of Neurological Surgery, Northwestern University, Chicago IL 60611, USA; Department of Physical Medicine & Rehabilitation, Northwestern University, Chicago IL 60611, USA; Shirley Ryan AbilityLab, Chicago, IL 60611, USA; Department of Neuroscience, Northwestern University, Chicago, IL 60611, USA; Department of Biomedical Engineering, Northwestern University, Evanston, IL 60201, USA

**Keywords:** LFPs, ECoG, phase-amplitude coupling, gamma, movement, speech

## Abstract

Planning and executing motor behaviors requires coordinated neural activity among multiple cortical and subcortical regions of the brain. Phase-amplitude coupling between the high-gamma band amplitude and the phase of low frequency oscillations (theta, alpha, beta) has been proposed to reflect neural communication, as has synchronization of low-gamma oscillations. However, coupling between low-gamma and high-gamma bands has not been investigated. Here, we measured phase-amplitude coupling between low- and high-gamma in monkeys performing a reaching task and in humans either performing finger movements or speaking words aloud. We found significant coupling between low-gamma phase and high-gamma amplitude in multiple sensorimotor and premotor cortices of both species during all tasks. This coupling modulated with the onset of movement. These findings suggest that interactions between the low and high gamma bands are markers of network dynamics related to movement and speech generation.

## Introduction

Local field potentials (LFPs) are generated largely by the ensemble postsynaptic activity of populations of neurons and reflect underlying network dynamics(*1*). Traditionally categorized into several canonical frequency bands, modulation of activity in the theta (*θ*, 4-8 Hz), mu/alpha (*µ*/*α*, 8-13 Hz), beta (*β*, 13-30 Hz), and gamma (*γ*, 40-200 Hz) bands is linked to a wide range of brain functions, such as memory(*2*), attention(*2, 3*), spatial navigation(*4*), language perception and production(*5-7*), movement and force production(*8-16*). Moreover, studies have demonstrated correlations between LFP rhythms and neuronal spiking(*14, 17-19*) and between LFP rhythms of different frequencies(*20, 21*), the latter case being termed cross-frequency coupling (CFC).

The rhythmicity of LFP oscillations offers an elegant potential mechanism for coordinating neural activity over a wide range of spatial and temporal scales; thus, LFPs are hypothesized by some to have functional roles by influencing neural activity(*10, 22-26*). In particular, the *γ* band has received substantial attention due to consistent observations of event-related modulations in *γ* band activity and synchronization over a wide range of behaviors and cortical regions(*12, 27, 28*). Although variably defined in the literature(*29*), in neocortex, the *γ* band is really two distinct bands, low gamma (L*γ*, variably defined but here defined as 40-50 Hz) and high gamma (H*γ*, 70-200 Hz), thought to represent different neural processes(*12, 14, 20, 30-33*). Indeed, oscillations mostly within the L*γ* band are theorized to have an important role in neural communication(*22, 27*), whereas H*γ* activity is broadband activity, traditionally considered a proxy for ensemble spiking activity(*31-33*).

The execution of motor behaviors involves coordinated neural activity within higher-order motor cortices (premotor and posterior parietal areas), subcortical nuclei and cerebellum, and the sensorimotor cortex, a region comprised by the primary motor (M1) and primary somatosensory (S1) cortices located on the pre- and postcentral gyri, respectively. Hypothesized as a marker of, and (by some) potentially the mechanism underlying, coordinated neural activity and information transfer within and between cortical networks(*34*), CFC describes the interactions between LFPs of different frequencies. One well-studied type of CFC is phase-amplitude coupling (PAC), in which the amplitude of higher frequency activity varies with the phase of a lower frequency rhythm. Many methods have been developed(*20, 35, 36*) and used to detect PAC between several different frequency band pairs during motor behaviors in both healthy and pathological states(*14, 20, 37-39*). For example, one seminal study reported coupling between *θ* phase and H*γ* amplitude (*θ*-H*γ*) over a range of sensorimotor and cognitive tasks across the human cortex(*20*). Additionally, several studies have observed *θ*-L*γ* and *θ*-H*γ* PAC in M1 of rats(*14*), *µ*/*α*-H*γ* PAC in the sensorimotor cortex of humans(*38*), and *β*-H*γ* PAC in the sensorimotor cortex of humans(*37, 39*) during motor behaviors. Some of these studies noted a decrease in *µ*/*α*-H*γ* and *β*-H*γ* PAC at movement onset, suggesting that these two types of PAC might suppress or “gate” movement(*37, 38*). One study also found L*γ*-spike coupling(*14*), suggesting that there might also be coupling between L*γ* and H*γ*. Moreover, PAC in the motor cortex and basal ganglia has been investigated as a biomarker to help control closed-loop deep brain stimulation in treatment of movement disorders(*39-45*).

Here we describe a novel form of PAC between L*γ* and H*γ* across species and motor behaviors. We recorded LFPs in monkeys and humans during three different motor behaviors of varying complexity: reaching, finger flexion, and speaking. We computed L*γ*-H*γ* PAC using two different methods: the modulation index (MI)(*35*) and a generalized linear model (GLM) framework(*36*). We found L*γ*-H*γ* PAC in many parts of the motor and premotor cortices of monkeys and humans and show that it modulates during motor behaviors. To our knowledge, this is the first study to investigate L*γ*-H*γ* PAC. This finding provides new insight into the roles of different gamma band activities in motor and premotor cortices. Additionally, L*γ*-H*γ* PAC could potentially serve as a biomarker for studies of motor control or movement disorders.

## Results

We collected intracranial recordings in monkeys and humans performing different motor behaviors (Figures 1a-1c). Two rhesus monkeys (C and M) performed a center-out reaching task with visual feedback, during which we recorded neural activity from two intracortical arrays in the primary motor cortices (M1) of both monkeys (CM1 and MM1) and from one array in the primary somatosensory cortex (S1) of one (MS1) over multiple experimental sessions spanning 4-9 weeks (Figure 1a). Furthermore, 12 human participants performed either a finger-flexion task (5 participants, Figure 1b) or a word-reading task (7 participants, Figure 1c, see Supplementary Table 1 for demographics). In these participants, we recorded neural activity from electrocorticography (ECoG) arrays covering the posterior frontal lobe and postcentral gyrus (see Methods for details).

**Figure 1.**
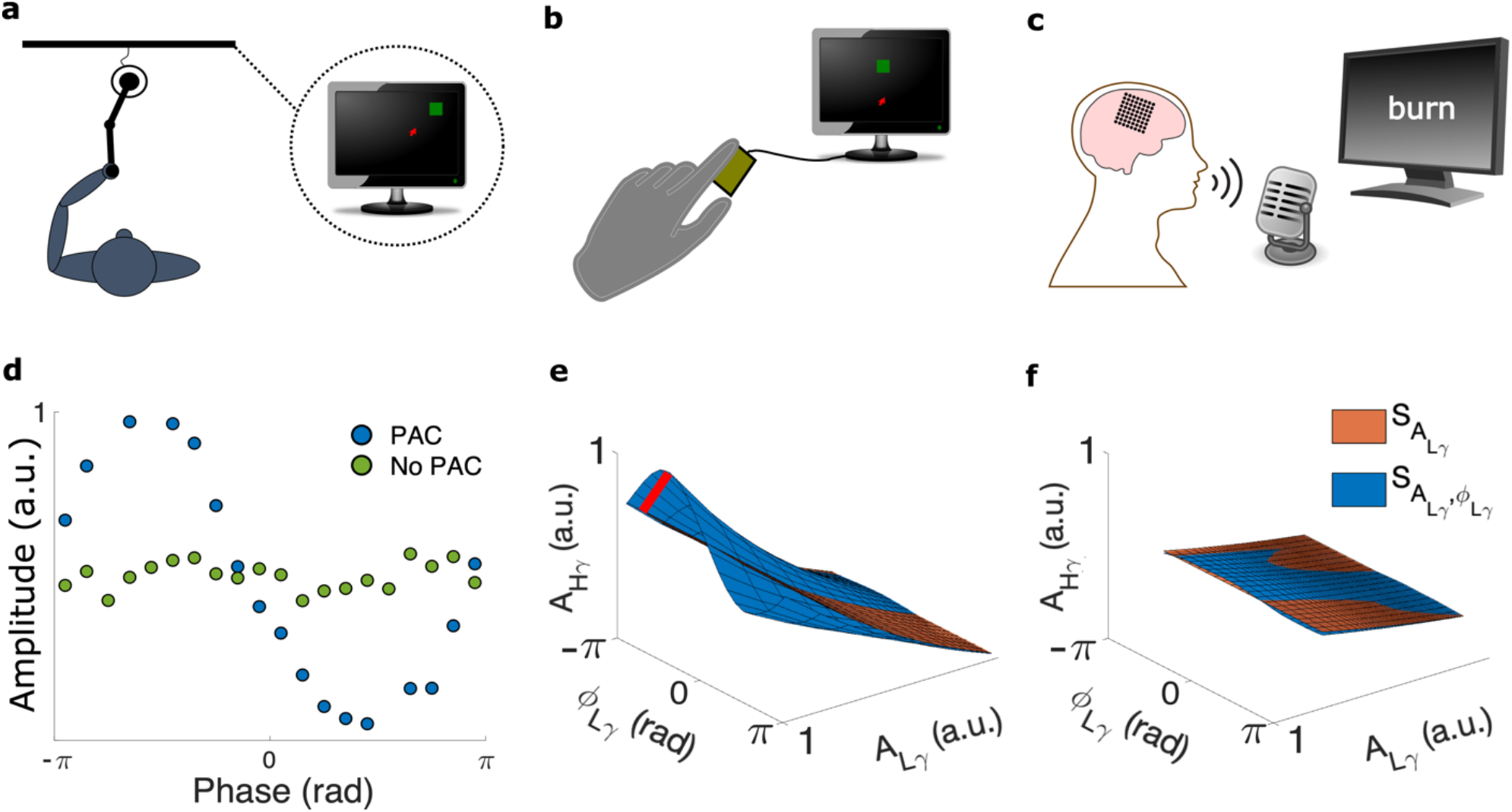
Motor behaviors and the phase-amplitude coupling (PAC) methods. **a** Monkey performing a center-out reaching task using a planar manipulandum. **b** Human participant performing a finger movement task. The participant flexed the index finger and used isometric force to move a computer cursor in 1D to randomly placed target (see Methods). **c** Human participant reading single words from the screen (word reading task). **d-f** Plots generated from an example electrode from the center-out task. **d** Example phase-amplitude histograms of H*γ* amplitude at each L*γ* phase. Notable variation of the higher frequency amplitude with phase indicates PAC (blue), whereas little-to-no variation suggests no PAC (green). a.u., arbitrary units. **e**,**f** Surfaces generated by the generalized linear model (GLM) framework demonstrating PAC (**e**) and no PAC (**f**). The orange surface (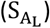) depicts the higher frequency amplitude (A_H_) as a function of the lower frequency amplitude (A_L_). The blue surface is the A_H_ as a function of both the lower frequency phase (ϕ _L_) and A_L_. The degree of PAC is directly proportional to the maximum orthogonal distance between the two surfaces (red line).

### L*γ*-H*γ* phase-amplitude coupling is a marker of rest and reaching in monkey M1

In the sensorimotor cortex, descriptions of *θ*-L*γ*(*14*), *µ*/*α*-H*γ*(*38*), and *β*-H*γ*(*37, 39*) PAC and their modulation by movement(*37, 38*) have provided insight into the temporal gating of motor representation in the sensorimotor cortex during movement execution. To add to these previous findings, we investigated the existence of L*γ*-H*γ* PAC in the sensorimotor cortex of monkeys and whether it modulates with movement. For each experimental session, we aligned the trials to the outward reach onset, seeing typical modulation of H*γ* power around reach onset in precentral and postcentral gyri (Figures 2a and 2b). We defined two intervals: resting baseline in the center target (−500 to -300 ms) and reach onset (−100 to 100 ms). For each electrode and interval, we estimated the L*γ*-H*γ* PAC using z-scored modulation index (MI_z_, Figure 1d)(*35*), and using a slightly modified version of a newer GLM framework (R_PAC_, Figure 1e)(*36*). We selected the second method because it considers the power of the frequency band defining phase when estimating PAC, reducing the impact of an important confound and thus permitting a more valid interpretation of PAC changes with behavior(*36, 46*). We only included electrodes demonstrating significant L*γ*-H*γ* PAC identified during either interval for statistical comparisons.

**Figure 2.**
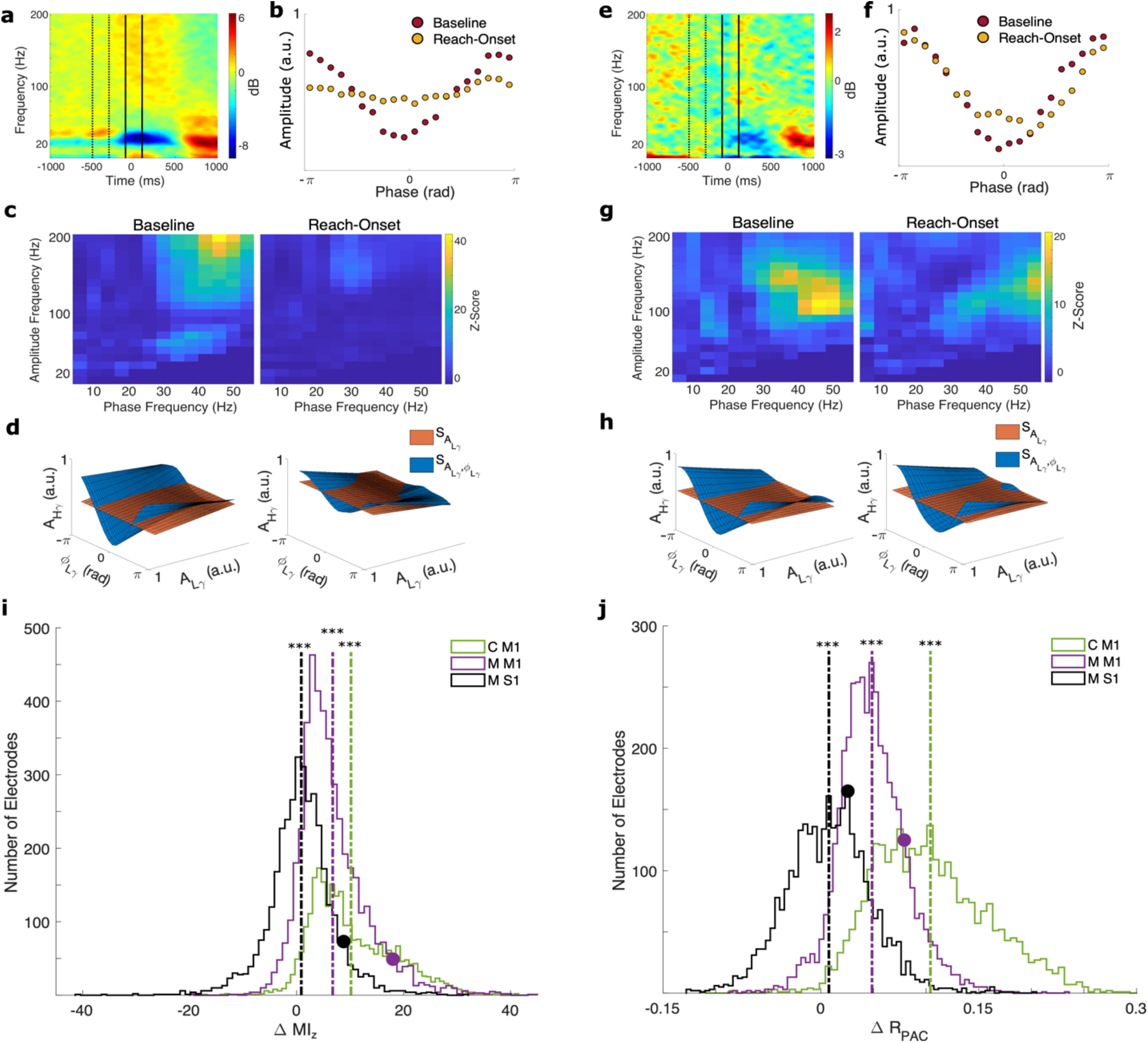
Low *γ*-high *γ* PAC in monkeys performing the center-out reaching task. **a-d** Illustrative plots from an example electrode and experimental session in M1 in monkey M (M M1). **a** Spectrogram time-locked to reach-onset with the baseline (−500 to -300 ms, dashed lines) and reach-onset (−100 to 100 ms, solid lines) intervals marked. **b** Phase-amplitude plots during the baseline (red) and reach-onset (yellow) intervals. **c** Comodulograms during the baseline (left) and reach-onset (right) intervals. **d** Surfaces generated by the GLM framework during the baseline (left) and reach-onset (right) intervals. The greater the differences between the surfaces, the greater the PAC. Thus, PAC decreases from baseline to reach onset. **e-h** Same as in a-d, except in monkey M S1 (M S1). **i** Distributions of the differences in the z-scored modulation index (∆MI_z_) between intervals (baseline minus reach-onset) in each electrode over all experimental sessions for M1 in monkey C (C M1), M M1, and M S1. Vertical dashed lines represent mean ∆MI_z_ for each monkey/region. Circles represent exemplary electrodes shown in **a**-**d** (purple) and **e-h** (black). ∆MI_z_ was significantly greater than 0 in all three cases (***, p<0.001), with much higher means in M1 and M1 than S1. **j** Same as in **i**, but for ΔR_PAC_ ΔR_PAC_ was significantly greater than 0 in all three cases, with much higher means in M1 than in S1.

We observed a high degree of L*γ*-H*γ* PAC in all three intracortical arrays using both the MI_z_ and GLM framework (Figure 2). Using the MI_z_, many electrodes demonstrated significant L*γ*-H*γ* PAC during either interval for CM1 (2728 of 2942, 92.7%), MM1 (4548 of 5435, 83.7%), MS1 (3633 of 4351, 83.5%). Similarly, using the GLM framework, many electrodes showed significant L*γ*-H*γ* PAC during either interval for CM1 (2899 of 2942, 98.5%), MM1 (4188 of 5435, 77.1%), and MS1 (3194 of 4351, 73.4%). For several electrodes in the preCG and postCG, comodulograms created using the MI_z_ demonstrated that L*γ*-H*γ* PAC was the predominant type of PAC (based on visual inspection), especially during the baseline interval (Figures 2c and 2g). This was not a consistent observation, as other types of PAC previously reported in the sensorimotor cortex, such as beta-H*γ* PAC(*37, 44*), were predominant in other electrodes (Supplementary Figure 1a).

We also found a differential modulation of L*γ*-H*γ* PAC with reaching based on brain region. Across all electrodes with significant L*γ*-H*γ* PAC identified with the MI_z_, the MI_z_ was significantly higher during baseline than during reach-onset in CM1 (two-tailed paired t-tests: t_2727_ = 67.70, p < 0.0001) and MM1 (t_4547_ = 69.37, p < 0.0001), and this pattern was seen in most electrodes in CM1 (2619 of 2728, 96.0%) and MM1 (4159 of 4548, 91.4%; Figure 2i). Likewise, across all electrodes with significant L*γ*-H*γ* PAC identified with the GLM framework, the baseline R_PAC_ was significantly higher than the reach-onset R_PAC_ in CM1 (two-tailed paired t-tests: t_2898_ = 102.81, p < 0.0001) and MM1 (t_4187_ = 97.00, p < 0.0001), and this pattern was seen in nearly all electrodes in CM1 (2875 of 2899, 99.2%) and MM1 (3978 of 4188, 95.0%; Figure 2j).

For MS1, we also observed significantly higher baseline L*γ*-H*γ* PAC than reach-onset L*γ*-H*γ* PAC using both MI_z_ (two-tailed paired t-tests: t_3632_ = 8.58, p < 0.0001) and R_PAC_ (t_3193_ = 10.66, p < 0.0001). However, the mean difference (Δ) in MI_z_ and R_PAC_ between the two intervals (i.e., baseline minus reach-onset) across all electrodes with significant L*γ*-H*γ* PAC was much smaller in MS1 (mean ΔMI_z_ = 0.87; mean ΔR_PAC_ = 0.0076) than in MM1 (mean ΔMI_z_ = 6.74; mean ΔR_PAC_ = 0.049; dashed lines in Figures 2i-j) and CM1 (mean ΔMI_z_ = 10.16; mean ΔR_PAC_ = 0.10; dashed lines in Figures 2i-j). Compared to the overall distribution of ΔMI_z_ in MS1, the distributions of ΔMI_z_ were significantly greater in both MM1 (two-tailed unpaired t-tests: t_8179_ = 41.48, p < 0.0001) and CM1 (t_6539_ = 53.09, p < 0.0001). Likewise, the distributions of ΔR_PAC_ in MM1(t_7380_ = 48.36, p < 0.0001) and CM1 (t_6091_ = 78.99, p < 0.0001) were significantly greater than the ΔR_PAC_ distribution in MS1. Indeed, a relatively smaller proportion of MS1 electrodes had a greater baseline than reach-onset MI_z_ (2082 of 3633, 57.3%) and R_PAC_ (1851 of 3194, 58.0%) compared to MM1 (MI_z_: 91.4%, R_PAC_: 95.0%) and CM1 (MI_z_: 96.0%, R_PAC_: 99.2%). These results indicate a regional influence on degree of modulation of L*γ*-H*γ* PAC with movement.

### L*γ*-H*γ* phase-amplitude coupling is a marker of finger flexion vs. rest in humans

Our initial results confirm the existence of L*γ*-H*γ* PAC in the sensorimotor cortex of monkeys and indicate a movement- and region-related modulation of L*γ*-H*γ* PAC. To support and expand upon these initial findings, we investigated L*γ*-H*γ* PAC in humans performing a finger-flexion task. Briefly, the participants were visually cued to flex their index finger, then extend back to a neutral baseline (see Methods). We categorized electrodes as precentral gyrus (preCG, including M1 and part of premotor cortex), postcentral gyrus (postCG), or anterior to the precentral sulcus region (aPreCS, including premotor and prefrontal cortices) electrodes depending on their estimated location. We defined baseline (−600 to -400 ms) and flexion-onset (−200 to 0 ms) intervals relative to the onset of finger flexion using slightly earlier times than for the monkeys because there were multiple electrodes in premotor cortex (which activates sooner) included in the analysis. We computed the MI_z_ and R_PAC_ for each interval and electrode, pooled results over all participants based on their respective interval and electrode category, and only included electrodes demonstrating significant L*γ*-H*γ* PAC during either interval for statistical comparisons.

We found L*γ*-H*γ* PAC in all three defined brain regions using both the MI_z_ and GLM framework (Figure 3). Using the MI_z_, we identified many electrodes in the preCG (24 of 98, 24.5%), postCG (6 of 26, 23.1%), and aPreCS (32 of 134, 23.9%) with significant L*γ*-H*γ* PAC during either interval. Of these electrodes, most in the preCG (18 of 24, 75.0%), postCG (4 of 6, 66.7%), and aPreCS (19 of 32, 59.4%) had a greater MI_z_ at baseline than flexion-onset (Figures 3f-g). Additionally, L*γ*-H*γ* PAC was the predominant or codominant type of PAC (based on visual inspection) in some electrodes but not all (Figure 3c and Supplementary Figure 1b). Using the GLM framework, we identified several electrodes in the preCG (25 of 98, 25.5%), postCG (1 of 26, 3.8%), and aPreCS (30 of 134, 22.4%) with significant L*γ*-H*γ* PAC during either interval. Of these electrodes, most in the preCG (17 of 25, 68.0%), postCG (1 of 1, 100%), and aPreCS (23 of 30, 76.7%) had a greater R_PAC_ at baseline than flexion-onset (Figures 3h-i).

**Figure 3.**
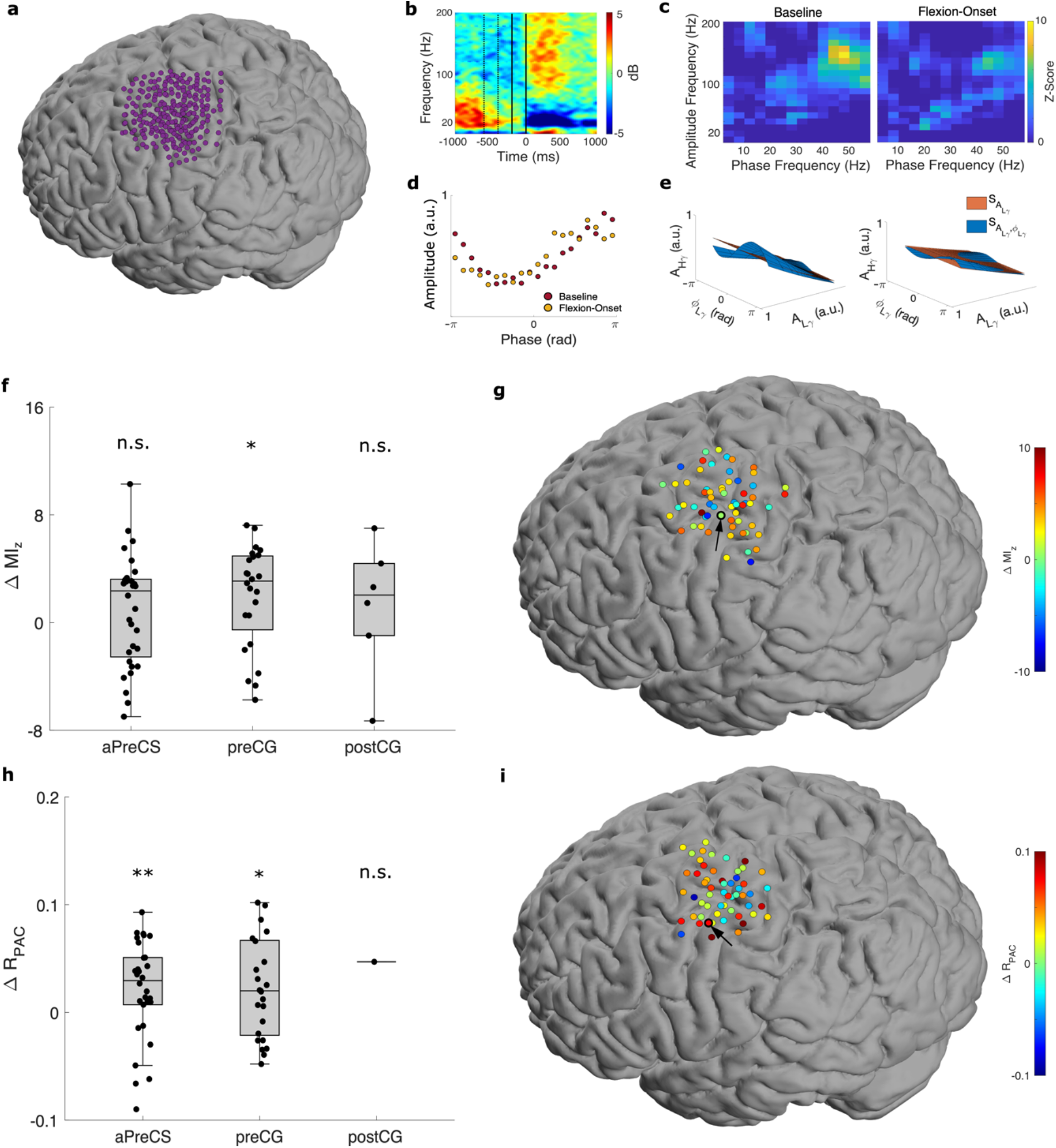
Low *γ*-high *γ* PAC in humans performing finger movements. **a** Spatial distribution of all electrodes across all five participants plotted on a template brain. **b-d** Illustrative plots from an example electrode in motor cortex (circled in black/arrow in **g**). **b** Spectrogram time-locked to flexion-onset (top, left) with baseline (−600 to -400 ms, dashed lines) and flexion-onset (−200 to 0 ms, solid lines) intervals marked. **c** Comodulograms during the baseline (left) and flexion-onset (right) intervals. **d** Phase-amplitude plots during the baseline (red) and flexion-onset (yellow) intervals. **e** Surfaces generated by the GLM framework during the baseline (left) and flexion-onset (right) intervals from another example electrode (circled in black/arrow in **i**). **f D**ifferences in MI_z_ between intervals (baseline minus flexion-onset) per electrode over all participants. ∆MI_z_ was significantly greater than 0 for precentral gyrus (preCG) electrodes (*, p<0.05), but not for anterior to the precentral sulcus (aPreCS) and postcentral gyrus (postCG) electrodes (n.s.). **g** Spatial distribution of ∆MI_z_ for significant electrodes with the example electrode marked (black). **h** Same as in c, except for ΔR_PAC_ ΔR_PAC_ was significantly greater than 0 for aPreCS (** p<0.01) and preCG electrodes (* p<0.05), but not for postCG electrodes (n.s.). **i** Same as in g, except for ΔR_PAC_

In monkeys, we showed that L*γ*-H*γ* PAC modulates with movement greatly in M1 and less so in S1 (Figures 2i and 2j). One advantage of ECoG over intracortical arrays is much broader coverage, allowing us to investigate L*γ*-H*γ* PAC in more areas. We computed the difference in the pooled MI_z_ and R_PAC_ (across patients and electrodes) between the two intervals (baseline minus flexion-onset) per cortical region. We found that in preCG, PAC significantly decreased moving from the baseline to flexion-onset interval using both pooled MI_z_ (median ΔMI_z_ = 3.08; one-tailed Wilcoxon signed rank test: W^+^ = 226, p = 0.015) and R_PAC_ (median ΔR_PAC_ = 0.020; W^+^ = 235, p = 0.026). In contrast, in postCG, there was no change between baseline and flexion-onset intervals in either the pooled MI_z_ (median ΔMI_z_ = 2.04; W^+^ = 14, p = 0.28) or R_PAC_ (ΔR_PAC_ = 0.047; W^+^ = 1, p = 0.50; Figure 3f and 3h). Interestingly, we observed no change in the pooled aPreCS MI_z_ (median ΔMI_z_ = 2.35; one-tailed Wilcoxon signed rank test: W^+^ = 323, p = 0.139) but did find a significant decrease in the pooled aPreCS R_PAC_ (median ΔR_PAC_ = 0.030; W^+^ = 350, p = 7.27e-3) moving from the baseline to flexion-onset interval (Figures 3f and 3h). Although the MI_z_ results showed only a nonsignificant trend, the significant decrease in R_PAC_ could indicate that the movement-related modulation of L*γ*-H*γ* PAC extends into the aPreCS, as the GLM framework permits a more accurate interpretation of PAC(*36*).

### L*γ*-H*γ* phase-amplitude coupling discriminates between silence and speech onset in humans

Thus far, we have demonstrated that L*γ*-H*γ* PAC and its modulation patterns with simpler movements are consistently present and generalize across species. To examine whether these patterns were present in other types of movement, we investigated L*γ*-H*γ* PAC in humans performing a more complex motor behavior—speech. We categorized electrodes as preCG, postCG, or posterior inferior frontal gyrus (pIFG) depending on their estimated location (Figure 4a). As in the finger flexion participants, we computed MI_z_ and RPAC for each categorized electrode during the baseline (−600 to -400 ms) and voice-onset (−200 to 0 ms) interval.

**Figure 4.**
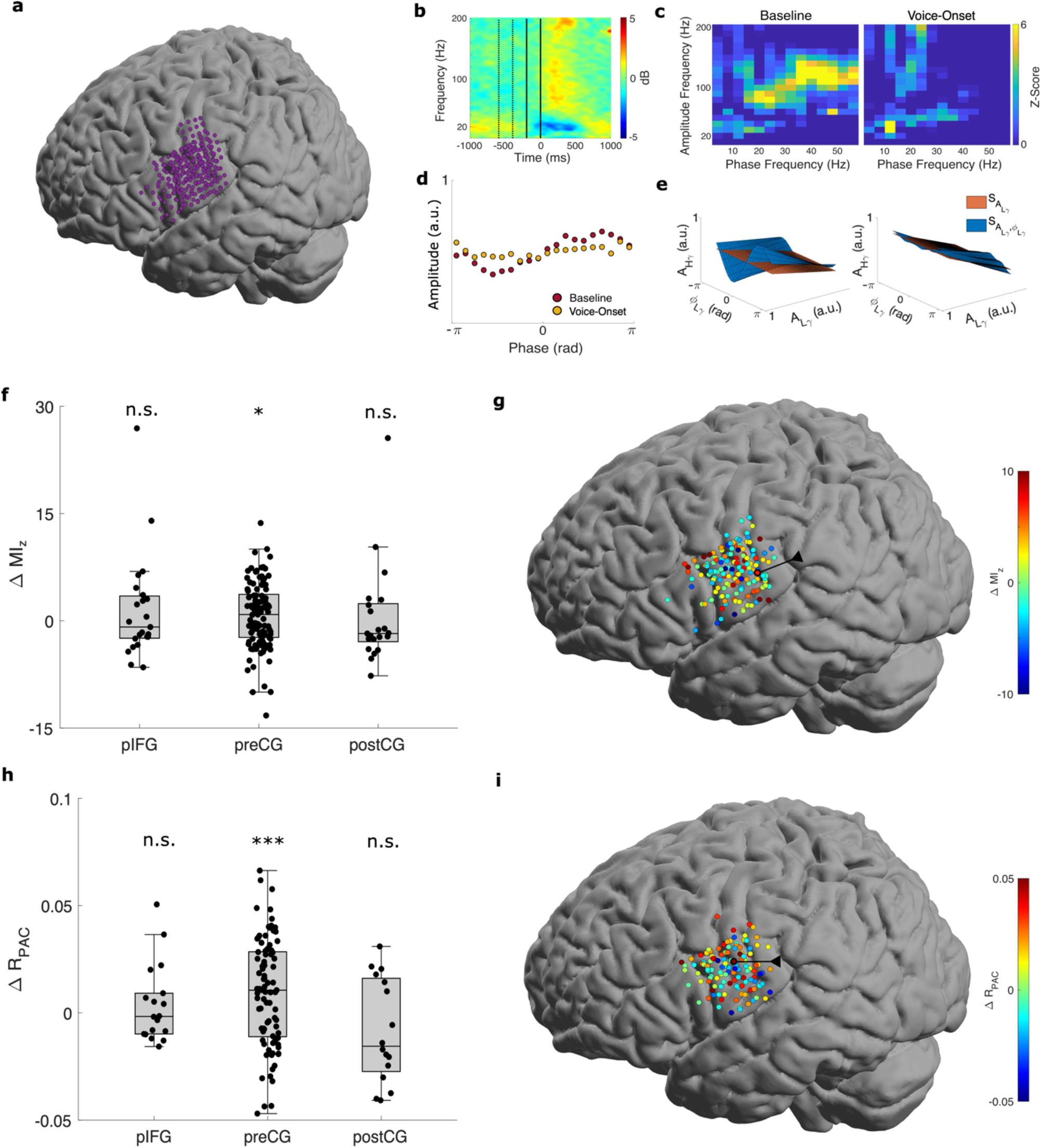
Low *γ*-high *γ* PAC in humans reading words. **a** Spatial distribution of all electrodes across all seven participants on a template brain. **b-e** Illustrative plots from an example electrode (circled in black in **g**). **b** Spectrogram time-locked to voice-onset (top, left) with baseline (−600 to -400 ms, dashed lines) and voice-onset (− 200 to 0 ms, solid lines) intervals marked. **c** Comodulograms during the baseline (left) and voice-onset (right) intervals. **d** Phase-amplitude plots during the baseline (red) and voice-onset (yellow) intervals. **e** Surfaces generated by the GLM framework during the baseline (left) and voice-onset (right) intervals from another example electrode (circled in black in **i**). **f** Differences in MI_z_ between intervals (baseline minus voice-onset) per electrode over all participants. ΔMI_z_ was significantly greater than 0 for preCG electrodes (* p<0.05), but not for posterior inferior frontal gyrus (pIFG) and postCG electrodes (n.s.). **g** Spatial distribution of ∆MI_z_ for significant electrodes with the example electrode marked (white). **h** Same as in c, except for ΔR_PAC_ ΔR_’#(_ was significantly greater than 0 for preCG electrodes (***), but not for pIFG (n.s) and postCG electrodes (n.s.). **i** Same as in g, except for ΔR_PAC_ (example electrode marked as black).

We again found L*γ*-H*γ* PAC in all three speech-related brain regions using both the MI_z_ and GLM framework (Figure 4). Using the MI_z_, we identified many electrodes in the preCG (109 of 172, 63.4%), postCG (21 of 44, 47.7%), and pIFG (23 of 48, 47.9%) with significant L*γ*-H*γ* PAC during either interval (Figures 4b-c). Of these electrodes, most in the preCG (58 of 109, 53.2%) and some in the postCG (7 of 21, 33.3%) and pIFG (10 of 23, 43.5%) had a greater baseline than voice-onset MI_z_ (Figures 4f-g). L*γ*-H*γ* PAC was the predominant or codominant type of PAC in some electrodes but not all (Figure 4c and Supplementary Figure 1c). Similarly, we identified many electrodes with significant L*γ*-H*γ* PAC during either interval using the GLM framework in the preCG (89 of 172, 51.7%), postCG (16 of 44, 36.4%), and pIFG (18 of 48, 37.5%). Of these electrodes, most in the preCG (56 of 89, 62.9%) and many in the postCG (6 of 16, 37.5%) and pIFG (8 of 18, 44.4%) had a greater baseline than voice-onset R_PAC_ (Figure 4h-i).

As we found for monkeys and humans doing finger movements, we found region-related modulation of L*γ*-H*γ* PAC around word vocalization (Figures 4f-i). The pooled preCG MI_z_ and R_PAC_ significantly decreased moving from the baseline to voice-onset interval (median ΔMI_z_ = 0.88; one-tailed Wilcoxon signed rank test: W^+^ = 3550, p = 0.048; median ΔR_PAC_ = 0.011; W^+^ = 2792, p = 6.23E-4). In contrast, the pooled MI_z_ and R_PAC_ in the pIFG (median ΔMI_z_ = -0.87; W^+^ = 150, p = 0.36; median ΔR_PAC_ = -0.0017; W^+^ = 90, p = 0.43) and postCG (median ΔMI_z_ = - 1.78; W^+^ = 95, p = 0.77; ΔR_PAC_ = -0.016; W^+^ = 43, p = 0.91) did not change between the two intervals (Figures 4f and 4h).

## Discussion

Generating movement and speech requires the coordination and control of neurons within brain motor and speech networks. Here, we examined cortical recordings in monkeys and humans for L*γ*-H*γ* PAC during and before movement and speech. We confirmed that L*γ*-H*γ* PAC is widespread across different motor regions, behaviors, and species. Furthermore, we observed a consistent, region-related modulation of L*γ*-H*γ* PAC during these motor behaviors across species. We found that L*γ*-H*γ* PAC was high in non-movement states and decreased at the onset of movement in both monkeys and humans in M1 and preCG, respectfully. These modulations were independent of L*γ* amplitude modulations. L*γ*-H*γ* PAC was much less prevalent, and remained relatively unchanged at movement onset, in S1 and postCG in monkeys and humans. Further, we observed similar, though less consistent, decreases in L*γ*-H*γ* PAC in higher-order motor regions of humans at the onset of movement. Moreover, these patterns held for reaching, finger flexion, and vocalization. Collectively, these results suggest that modulation of L*γ*-H*γ* PAC is a motor-related phenomenon indicative of underlying network dynamics fundamental to the activation of motor behaviors.

Event-related modulation of L*γ* and H*γ* activity has been observed in many brain regions, in several species, and during both motor and non-motor behaviors(*6, 12, 28, 33, 47*). Although sometimes combined in analyses, L*γ* and H*γ* are distinct entities associated with different origins and neural processes(*1, 12, 14, 20, 30-33*). L*γ* activity is thought to arise from rhythmic interactions between reciprocally connected inhibitory interneurons and excitatory pyramidal neurons(*48*). In contrast, H*γ* is thought to be broadband (non-oscillatory) activity likely arising from summed postsynaptic potentials of many thousands of neurons(*1, 37*), and as such is somewhat correlated with ensemble spiking activity(*31-33*). Functionally, observations of spike-L*γ* phase coupling(*14, 19, 23, 49-51*) and synchronization of L*γ* phase across brain areas led to the communication through coherence (CTC) hypothesis(*22, 27*), which posits that L*γ* band has a mechanistic role in neural communication by helping to synchronize across brain areas. Although the ability for L*γ* activity to directly influence neural activity is controversial(*25, 29, 52, 53*), it appears clearer that *γ* activity, especially in the L*γ* range, is at least a marker of engaged, cross-area neural networks(*33, 53*). For example, L*γ* activity may coordinate spiking between hippocampus and rhinal cortices(*54*), consistent with the observation that increasing L*γ* activity via biofeedback correlates with increased spiking synchronization(*53*). It also plays a strong role in spatial memory consolidation, as shown by causal closed-loop control of L*γ*(*55*). Moreover, in an Alzheimer disease mice model, optogenetic L*γ* stimulation restored previously diminished L*γ* activity and improved spatial memory(*56*).

L*γ* synchronization (CTC) and PAC are both thought to be indicative of information transfer in a cortical network(*22, 27, 34*). Moreover, L*γ* synchronization and PAC are related and may interact with each other(*57*). Yet, to our knowledge, this study is the first to extensively investigate and report the presence of interactions between L*γ* and H*γ* activity via PAC. While we cannot definitively assign a mechanistic role to L*γ*-H*γ* PAC due to limitations of PAC analysis(*46*), one interpretation of our results is that this phenomenon is a signature of a fundamental neural process that suppresses motor-related activity on a more local scale. This is similar to reports that *µ*/*α*-H*γ*(*38*) and *β*-H*γ*(*37*) PAC decrease with movement, suggesting an inverse relationship with (sometimes called gating of) motor activity. Accordingly, local release from this suppressive process occurs in areas important for generating the desired movement— such as regions of M1 projecting to agonist muscles—which is reflected by a decrease in L*γ*-H*γ* PAC in electrodes recording from these areas. On a larger spatial scale, a more global reduction in this suppressive process, reflected by a net decrease in L*γ*-H*γ* PAC over a region, permits the transition from an inactive to active motor state.

What is the neural process that gives rise to L*γ*-H*γ* PAC? Since H*γ* activity has been hypothesized to be a marker of ensemble spiking activity(*31-33*), one possibility is that L*γ*-H*γ* PAC is the LFP representation of spike-L*γ* correlative metrics, such as spike-L*γ* coherence. This would relate L*γ*-H*γ* PAC to the theorized functions of L*γ* activity(*22, 27*). Indeed, in M1 of rats performing forelimb movements, spiking activity in shallow cortical layers preferentially occurred at specific L*γ* phases(*14*). Since multiple animal studies have investigated spike-L*γ* correlations(*14, 19, 49, 51, 53, 58*), especially in a sensory context, it would be interesting to see if L*γ*-H*γ* PAC is also present in similar scenarios to support this possibility. If so, L*γ*-H*γ* PAC as a surrogate for spike-L*γ* correlative metrics could be a useful investigative tool, especially in humans. Spiking information is difficult to obtain in this population, and surface electrode arrays and depth electrodes provide opportunities to record neural activity across large spatial scales. Alternatively, H*γ* has been shown to be correlated with underlying latent spiking dynamics(*59*). It remains to be seen how L*γ*-H*γ* PAC may relate to the latent spiking dynamics.

L*γ* amplitude has been shown to decrease with movement in M1(*14*). Since modulations in the band activity defining phase can modulate PAC(*36, 46*), a simpler explanation for our findings is that the decrease in L*γ*-H*γ* PAC reflects decreased L*γ* amplitude. Although we cannot completely exclude this possibility, multiple pieces of evidence make it unlikely. Primarily, we utilized a modified GLM method that accounts for the amplitude of the band defining phase when estimating PAC strength, thus minimizing the effect of L*γ* amplitude on the estimated L*γ*-H*γ* PAC(*36*). While the modulation index method does not directly account for L*γ* amplitude, we observed an increase in the L*γ*-H*γ* MI_z_ with movement despite a corresponding decrease in L*γ* activity in some electrodes (Supplemental Figure S2a)(*60*). Additionally, in some electrodes, we observed little to no change in the L*γ*-H*γ* MI_z_ with movement despite a corresponding decrease in L*γ* activity (Supplemental Figure S2b). Moreover, in some electrodes with relatively strong L*γ* activity, we observed no significant L*γ*-H*γ* MI_z_ (Supplemental Figure S2c). Finally, in some electrodes, we observed decreases in L*γ*-H*γ* MI_z_ with movement despite little change in L*γ* activity (Supplemental figure S2d).

Beyond providing insight on network dynamics, PAC may also have practical applications. In patients with Parkinson’s disease (PD), there is exaggerated *β*-*γ* PAC in motor regions that reduce with therapeutic deep brain stimulation (DBS) of the subthalamic nucleus(*39, 44*). In addition to being a biomarker for PD, *β*-*γ* PAC has potential use as a feedback signal for closed-loop (adaptive) DBS(*39-45*). In epilepsy, *β*-H*γ* PAC has been used to detect seizures during invasive monitoring for epilepsy surgery(*61*). Furthermore, *θ*-L*γ* PAC may represent promising neurophysiological markers of schizophrenia and Alzheimer’s disease(*56, 62*). Additionally, several types of PAC, including L*γ*-H*γ* PAC, contain some information about speech that may be used for simple decoding tasks(*63*). These studies demonstrate possible investigational and translational avenues for L*γ*-H*γ* PAC.

## Methods

All experimental protocols were performed with approval from the Institutional Animal Care Use Committee and the Institutional Review Board of Northwestern University.

### Center-out task subjects and data acquisition

The monkey experimental protocols and results are reported in detail elsewhere(*64*). To summarize, two rhesus monkeys (C and M) were trained to perform a center-out reaching task while grasping a two-link planar manipulandum. The center-out task involved moving a computer cursor via the manipulandum to one of eight square, 2-cm outer targets spaced at 45° intervals around a circle of radius 10 cm. Each trial began with the monkey holding the cursor in the center target of the circle. After a random hold time of 0.5-0.6 s, a randomly selected outer target illuminated, signaling the monkey to reach to that target. The monkey needed to move the cursor into the outer target within 1.5 s and hold for a random time of 0.2-0.4 s to receive a liquid reward.

An intracortical 96-channel silicon microelectrode array (Blackrock Neurotech) was implanted in the proximal arm area of M1 contralateral to the tested arm in monkeys C and M. Another intracortical 96-channel array was previously implanted in the proximal arm area of S1 contralateral to the tested arm in monkey M. Intracortical arrays were grounded to the Cereport pedestal and referenced to a subdural platinum wire with 3 mm exposed length. Anesthesia and surgery details are described elsewhere(*64, 65*).

Neural signals were recorded using a 128-channel acquisition system (Cerebus, Blackrock Neurotech) while the monkeys performed the center-out task. LFPs were obtained by band-pass filtering between 0.5 and 500 Hz and sampling at 2 kHz for monkey C and 1 kHz for monkey M, with subsequent notch filtering at 60, 120, 180, and 240 Hz to remove line noise. Multiple data files of 5-20 min duration were recorded in each 60-90 min long experimental session. Overall, we analyzed 32 data files over 10 experimental sessions from C M1, 58 data files recorded over 11 sessions from M M1, and 48 data files recorded over 10 sessions from M S1. Movement onset was detected from synchronized kinematics recorded from the manipulandum as described in(*64, 65*).

### Finger movement task and data acquisition

All human participants were recruited at Northwestern Memorial Hospital and gave informed consent prior to participation. The experimental protocols and results are reported in detail elsewhere(*66*). Briefly, we analyzed recordings from five male human participants, four (FM1-FM4) undergoing awake intraoperative mapping prior to resection of low-grade gliomas and one (FM5) undergoing extraoperative intracranial monitoring before resection for medically refractory epilepsy. Participants were instructed to execute repeated trials of a finger movement task that required isotonic movement and isometric force of a single finger in sequence. At the beginning of each trial, participants held their index finger in a neutral posture. After a cue on a monitor, they executed a flexion movement, bringing the palmar surface of the distal phalanx of the index finger into contact with a load cell. They then applied force to match a randomly generated force target presented on the monitor within 2 s. Following a successful match or failure, the participant returned the finger to the neutral position. The next trial began after a delay of 1 s. Finger kinematics were recorded with a 22-sensor CyberGlove (Immersion), sampled at 2 kHz.

In FM1-4, ECoG arrays were over hand motor areas contralateral to the tested hand, which were defined using anatomical landmarks (i.e., “hand knob” in the precentral gyrus), preoperative fMRI and/or direct electrocortical stimulation mapping to identify functional hand motor area. In FM5, arrays were placed according to clinical necessity. All participants had arrays covering M1 and premotor cortex, with all except for FM3 covering S1 as well. For FM1-FM4, 64-electrode (8 × 8) higher-density arrays (Integra), with 1.5-mm exposed recording site diameter and 4-mm interelectrode spacing, were used. For FM5, a 32-electrode (8 × 4) array, with the same electrode size and spacing as the 64-electrode arrays, was used. ECoG signals were bandpass filtered from 0.3-500 Hz and sampled at 2 kHz, and force and kinematics were synchronously recorded, using a Neuroport Neural Signal Processor (Blackrock Microsystems).

### Word reading task and data acquisition

The experimental protocols and results are reported in detail elsewhere(*67, 68*). Briefly, we analyzed data from seven human participants, six (WR1-WR6) during awake intraoperative mapping for glioma resection and one (WR7) during extraoperative intracranial monitoring for medically refractory epilepsy. A monitor presented randomly selected, single words either every 2 s (WR1-6) or every 4 s (WR7). Participants read the word aloud as soon as it appeared. Words were monosyllabic, primarily with consonant-vowel-consonant (CVC) structure (details elsewhere(*69*)). Speech audio was recorded with microphones placed near the participant’s mouth and sampled at either 48 kHz (WR1-WR6 Califone) or 44.1 kHz (WR7).

ECoG arrays were placed over ventral M1 and ventral premotor cortex (PreCG) and frontal operculum (IFG). All electrode arrays except for WR6 covered portions of ventral S1 as well. Array location was confirmed as described above. Recordings in WR1-WR6 used 64-electrode, higher-density arrays and were recorded using the methods described above. Recordings in WR7 used a 32-electrode (8 × 4) clinical array (PMT), with 2.3-mm exposed diameter and 10-mm interelectrode spacing, and was recorded with a Nihon Kohden system, bandpass filtering from 0.5-300 Hz and sampling at 1 kHz. Audio recordings were synchronized to the ECoG recordings.

### ECoG electrode localization

For intraoperative recordings, electrode locations were stereotactically registered at the time of grid placement using Brainlab Curve. We identified each electrode’s functional anatomical position with regards to surrounding landmarks (i.e., central sulcus, precentral gyrus, frontal gyri) using the superposed electrode locations on the reconstructed cortical surface provided in the Brainlab software suite, as well as intraoperative photos. For extraoperative recordings, we used the Fieldtrip toolbox(*70*) to reconstruct the patients’ cortical surface from the pre-implantation MRI and co-registered it to the post-implantation CT scan. We verified our presumed electrode functional anatomical locations in both settings to be coherent with cortical stimulation mapping results. For ensemble visualization of electrodes from multiple participants, we translated our identified electrode positions to a template brain(*71*) using LeGUI software(*72*).

### Signal processing

All intracortical and ECoG signals were resampled to 1 kHz. All electrodes were visually inspected for noise or artifacts and excluded from subsequent analyses if noisy. The clean channels in each ECoG array were common average referenced (CAR). Trials were aligned to event onsets of each task. Movement onset in the reach and finger flexion tasks was computed from change in the 2D cursor velocity^62^ or finger position^64^. Speech onset was determined manually from the audio signal and spectrogram^66^. LFP and ECoG spectrograms were computed in a 2-s interval surrounding event onset using 256-ms bins of data, shifted in 25-ms increments. For each bin, a Hanning window and fast-Fourier transform was applied, and the resulting complex magnitudes were squared. Spectrograms were created by computing the log of the mean magnitude over trials for each bin and normalizing by subtracting the log of the mean power spectrum over the entire interval.

### Estimation of phase-amplitude coupling

Phase-amplitude coupling (PAC) between the phase of the lower frequency band and the amplitude of the higher frequency band was computed using two methods: the modulation index (MI)(*35*) and a modified generalized linear model (GLM) framework(*36*). For each method, the CAR signals were first bandpass filtered with a two-way least-squares FIR filter using EEGLAB’s *eegfilt*.*m*(*73*) to isolate activity within the L*γ* (40-50 Hz) and H*γ* (70-200 Hz) bands. We deliberately chose 40-50 Hz to represent L*γ* to avoid any potential overlap with beta and with line noise. Each band’s filtered signal was then z-scored in time. The Hilbert transform was then applied to these signals to extract the instantaneous L*γ* phase (*ϕ*_L*γ*_) and H*γ* amplitude (A_H*γ*_), as well as the instantaneous L*γ* amplitude (A _L*γ*_) for the modified GLM framework. Baseline and event-onset intervals were -500 to -300 ms and -100 to 100 ms, respectively, in monkeys and -600 to -400 ms and -200 to 0 ms, respectively, in humans. We chose slightly earlier intervals in humans because these recordings included premotor areas (anterior part of PCG and anterior to the PCG), which activate earlier than M1 and S1 for a given movement.

To estimate PAC for each interval using the MI(*35*), the corresponding 200 ms bins of *ϕ* _L*γ*_ and A_H*γ*_ from each trial were concatenated and sorted to create a histogram of amplitudes as a function of phases (20 phase bins equally spaced from -*π* to *π*). The MI value was then computed from the Kullback-Leibler divergence between the amplitude distribution as a function of *ϕ*_L*γ*_ and a uniform distribution. We then randomly shuffled trial pairs of *ϕ*_L*γ*_ and A_H*γ*_ 1000 times to create a distribution of surrogate MI values. The *z*-scored MI (MI_z_) was then computed by comparing the observed MI value to the mean MI value of the surrogate distribution, specifically

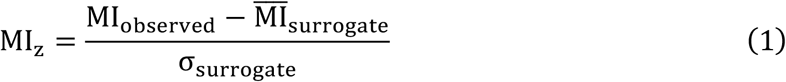

where higher values of MI_z_ suggest stronger PAC.

Comodulograms were created using the MI by first defining two sets of frequency bands, one for the phase frequencies and the other for the amplitude frequencies. The phase frequency bands were centered on frequencies ranging from 4 to 56 Hz in steps of 4 Hz and had fixed bandwidths of 4 Hz. The amplitude frequency bands were centered on frequencies ranging from 10 to 200 Hz in steps of 10 Hz and had variable bandwidths. Specifically, the bandwidths of the amplitude frequency bands were twice the center frequency of the corresponding phase band, as this ensured that the passband encompassed the sidebands created by the assumed phase frequency(*74*). The MI_z_ for each pair of phase and amplitude frequency bands was then computed as previously described to create comodulograms during the baseline and onset intervals.

To estimate PAC for each interval using the modified GLM framework(*36*), the corresponding 200-ms bins of *ϕ*_L*γ*_, A_H*γ*_, and A_L*γ*_ from each trial were concatenated and used to create three GLMs. Each GLM used a gamma distribution to model the conditional distribution of the response variable—A_H*γ*_—given the predictor variable, where the mean parameter of the gamma distribution *µ* was related to the predictor variable via a link function. The first GLM (GLM_1_) defined the link function as a linear combination of spline basis functions to approximate the predictor variable, *ϕ*_L*γ*_. The second GLM (GLM_2_) defined the link function as a linear function to approximate the predictor variable, A_L*γ*_. The third GLM (GLM_3_) defined the link function as a linear combination of the first two GLMs’ link functions and two terms that approximated the interaction between two predictor variables, *ϕ*_L*γ*_ and A_L*γ*_.

For PAC, GLM_2_ and GLM_3_ were used to create surfaces in the 3D space spanned by *ϕ*_L*γ*_ A_L*γ*_, and A_H*γ*_ (*36*). The surface created with GLM_2_, 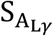 represented A_H*γ*_ as a function of only A_L*γ*_ and was thus constant in the *ϕ*_L*γ*_ dimension. The surface created with GLM_3_, 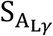 represented A_H*γ*_ as a function of both *ϕ*_L*γ*_ and A_Lγ._ The method’s measure of PAC, R_PAC_, was the maximum absolute fractional difference between these two surfaces, defined as

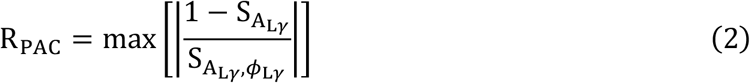

where higher values of R_PAC_ indicated stronger PAC. Surrogate R_PAC_ values were created by randomly shuffling the trial pairs of *ϕ*_L*γ*_ A_L*γ*_ and A_H*γ*_ and using the resulting concatenated signals to create the three GLMs. This was done 1000 times to create a surrogate distribution.

Analysis of phase-amplitude coupling

To identify electrodes with significant PAC within the baseline and onset intervals using the MI, the MI_z_ values during each interval were converted to one-sided *p*-values and corrected for the number of electrodes (false discovery rate correction, *α* = 0.05). To do the same using the modified GLM framework, *p* values were defined as the proportion of surrogate R_PAC_ values greater than the estimated R_PAC_ and corrected for the number of electrodes (false discovery rate correction, *α* = 0.05). If the proportion was 0, then *p* was set to 0.0005(*36*). Only electrodes with significant PAC during either the baseline or event onset intervals were included for further analysis. For the participants performing finger movements, each electrode was labeled as either a precentral gyrus (preCG), postcentral gyrus (postCG), or region anterior to the precentral sulcus electrode (aPreCS). For the participants reading words, each electrode was labeled as either a preCG, postCG, or posterior inferior frontal gyrus electrode.

For each interval during the reaching task, the degree of L*γ*-H*γ* PAC was determined by computing the proportion of electrodes with significant PAC per file. Differences in L*γ*-H*γ* PAC between the intervals was assessed by subtracting the MI_z_ and R_PAC_ during the onset interval from the MI_z_ and R_PAC_ during the baseline interval, respectively, for each included electrode across all files. For each defined brain region and each interval in human participants, the degree of L*γ*-H*γ* PAC was determined by computing the proportion of electrodes with significant PAC per participant. Differences in L*γ*-H*γ* PAC between the intervals was assessed by subtracting the MI_z_ and R_PAC_ during the onset interval from the MI_z_ and R_PAC_ during the baseline interval, respectively, for each included electrode in a region. Parametric and non-parametric statistical tests were used to compare PAC strength between intervals as specified in the Results.

## Acknowledgments

We thank Zachary Wright, Michael Scheid, and Luke Jordan for assistance collecting monkey data, and Stephan Schuele and our EEG technologists for assisting with recruitment and recording of ECoG data.

This research was supported in part by NIH grants K08NS060223 R01NS094748, R01 NS099210, R01NS112942, and F32-DC-015708 (to EMM); the Dixon Translational Research Grants Initiative at Northwestern Medicine and the Northwestern University Clinical and Translational Sciences Institute (NIH UL1RR025741, UL1-TR-000150 and UL1-TR-001422), Paralyzed Veterans of America Research Grant #2728, Brain Research Foundation (BRF SG 2009-14), Doris Duke Charitable Foundation Clinical Scientist Development Award #2011039, and a Craig H. Neilsen Foundation Fellowship (to RDF).

J.Z.N. wrote most of the code, analyzed the data, and wrote the manuscript. R.D.F. helped design the study, wrote some of the code, and collected some data. P.P. helped design the study and collected some data. J.K.H helped design the study and collected some data. E.M.M. collected some data. M.C.T. collected some data. J.M.R collected some data. M.W.S. is the Principal Investigator of this study.

The authors declare no conflicts of interest.

