## Supplementary Materials for "High-gamma activity is coupled to low-gamma oscillations in precentral cortices and modulates with movement and speech"

**Supplemental Table 1.** Participant demographics

| Participant | Sex | Age (years) |
|---|---|---|
| FM1 | M | 48 |
| FM2 | M | 33 |
| FM3 | M | 60 |
| FM4 | M | 30 |
| FM5 | M | 34 |
| WR1 | M | 57 |
| WR2 | M | 72 |
| WR3 | M | 43 |
| WR4 | M | 61 |
| WR5 | F | 45 |
| WR6 | M | 54 |
| WR7 | M | 50 |

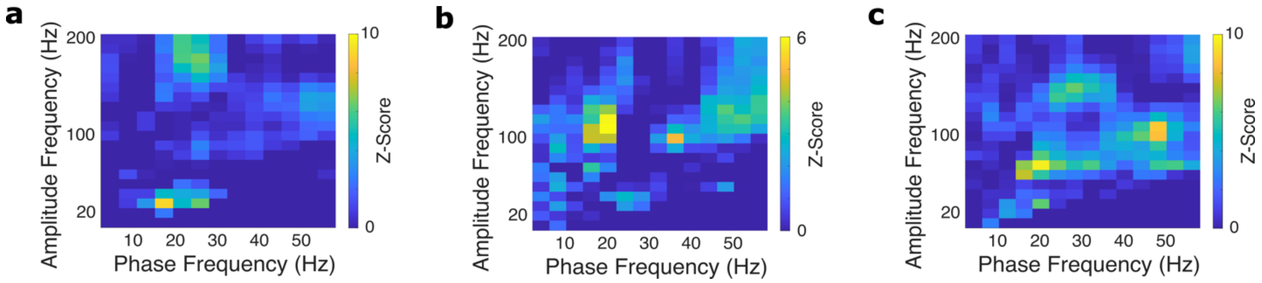


**Supplementary Figure 1.** Comodulograms of example electrodes demonstrating other types of phase-amplitude coupling in addition to low $\gamma$-high $\gamma$ PAC during the **a** center-out reaching task, **b** finger-movement task, and **c** word-reading task.

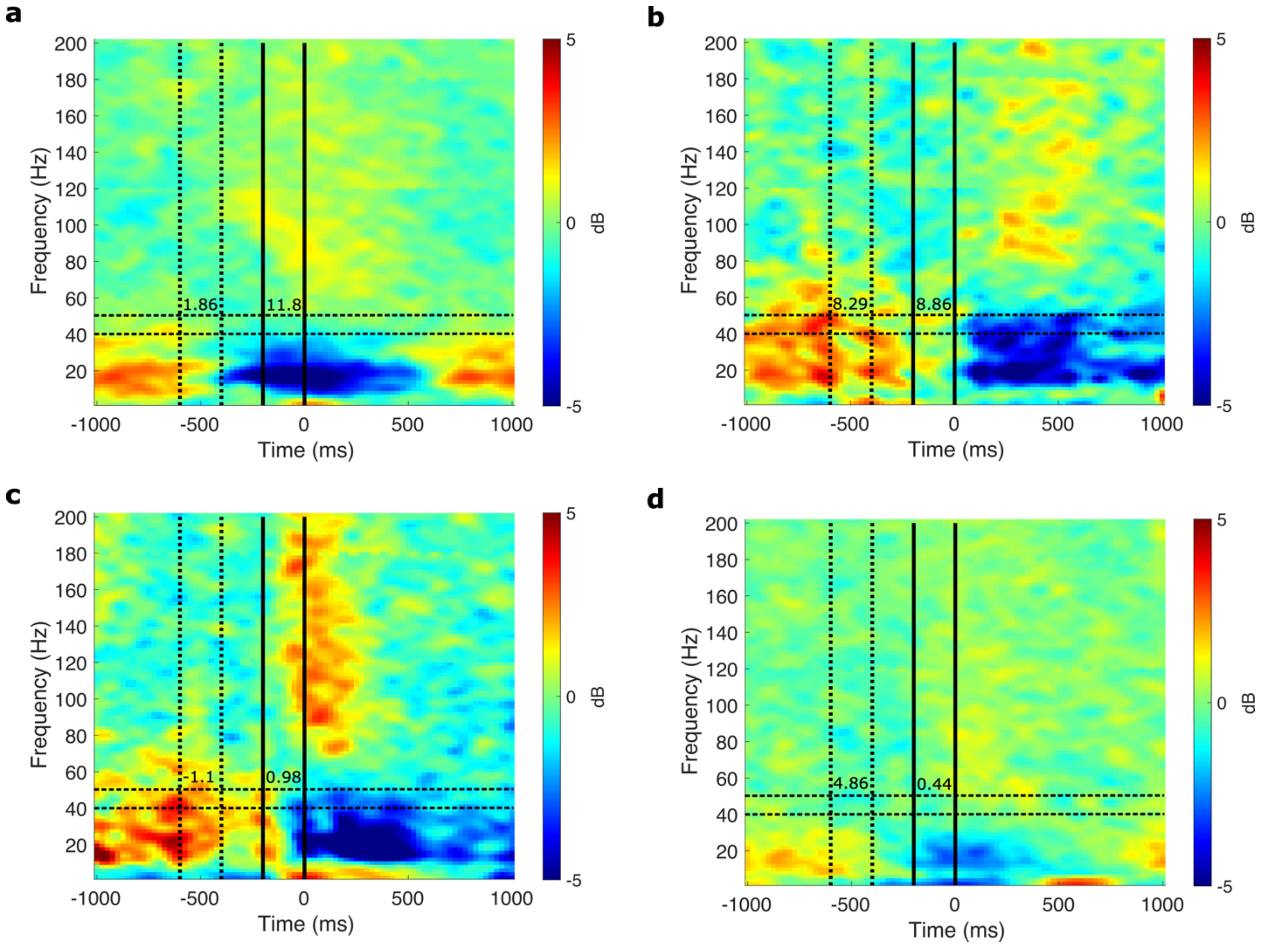


**Supplementary Figure 2.** Spectrograms of example ECoG electrodes during the finger-movement and word-reading tasks. **a** Decrease in L$\gamma$ (40-50 Hz, horizontal dashed black lines) power (by 0.80 dB) from the baseline (vertical dotted black lines) to motor-onset (vertical solid black lines) intervals in an electrode with significant L$\gamma$-H$\gamma$ PAC. PAC was measured by the z-scored modulation index ($MI_z$), during both intervals (specified by the numbers between the lines for each interval). The $\Delta MI_z$ between the two intervals (baseline minus motor-onset) was -9.94, indicating an increase in L$\gamma$-H$\gamma$ PAC with motor onset. **b** Decrease in L$\gamma$ power from the baseline to motor-onset intervals in an electrode with significant $MI_z$ during both intervals. $\Delta MI_z$ was -0.58, suggesting little to no change in L$\gamma$-H$\gamma$ PAC with motor-onset. **c** An electrode with no significant $MI_z$ during either interval despite relatively high L$\gamma$ power during baseline. **d** Small to no change in L$\gamma$ power (change of -0.26 dB) from baseline to motor-onset in an example electrode with significant $MI_z$ during the baseline interval that decreases substantially with motor onset.
